# Physiological alterations in microglial morphology associate with the sleep-wake cycle in a brain region-specific manner

**DOI:** 10.1101/2022.03.04.482976

**Authors:** Sarah Katharina Steffens, Tarja Helena Stenberg, Henna-Kaisa Wigren

## Abstract

Long-term total sleep deprivation induces changes in cortical - and hippocampal microglial morphology that closely resemble the microglial response to the gram-negative bacterial cell wall component lipopolysaccharide (LPS). A recent study found evidence that microglia could modify vigilance-states/sleep, but only few studies investigated microglial throughout the diurnal behavioral inactivity/activity pattern or the naturally occurring sleep-wake cycle, and those who have, only concentrated on the cortical or hippocampal microglia. As microglia demonstrate regional heterogeneity, we compared microglial diurnal morphological alterations in the somatosensory cortex (SC) and dorsal hippocampus (HC) to the basal forebrain (BF), which is a subcortical brain area involved in the regulation of vigilance states.

We collected mouse brain samples every 3h throughout the 24h light-dark-cycle and applied a 3D reconstruction method for the acquired confocal microscopy images for each brain area separately. While microglial regional heterogeneity was evident, stimulation of microglia with LPS caused comparable microglial responses in all brain areas. When comparing microglial features between the 12h light- and dark periods, regional heterogeneity re-appeared. As most of the morphological alterations occurred during the light period-the habitual sleeping period of the mice, we performed polysomnography to study the possible interaction of microglial morphology and sleep. We found that cortical-, but not HC- or BF microglial territory and volume negatively correlated with sleep slow wave activity (SWA), an electroencephalic feature of non-REM sleep (NREMS). Since microglia are sensitive to neuronal activity, we propose that the regional differences reflect vigilance-state specific neuronal activity patterns.

**Table of contents image:** 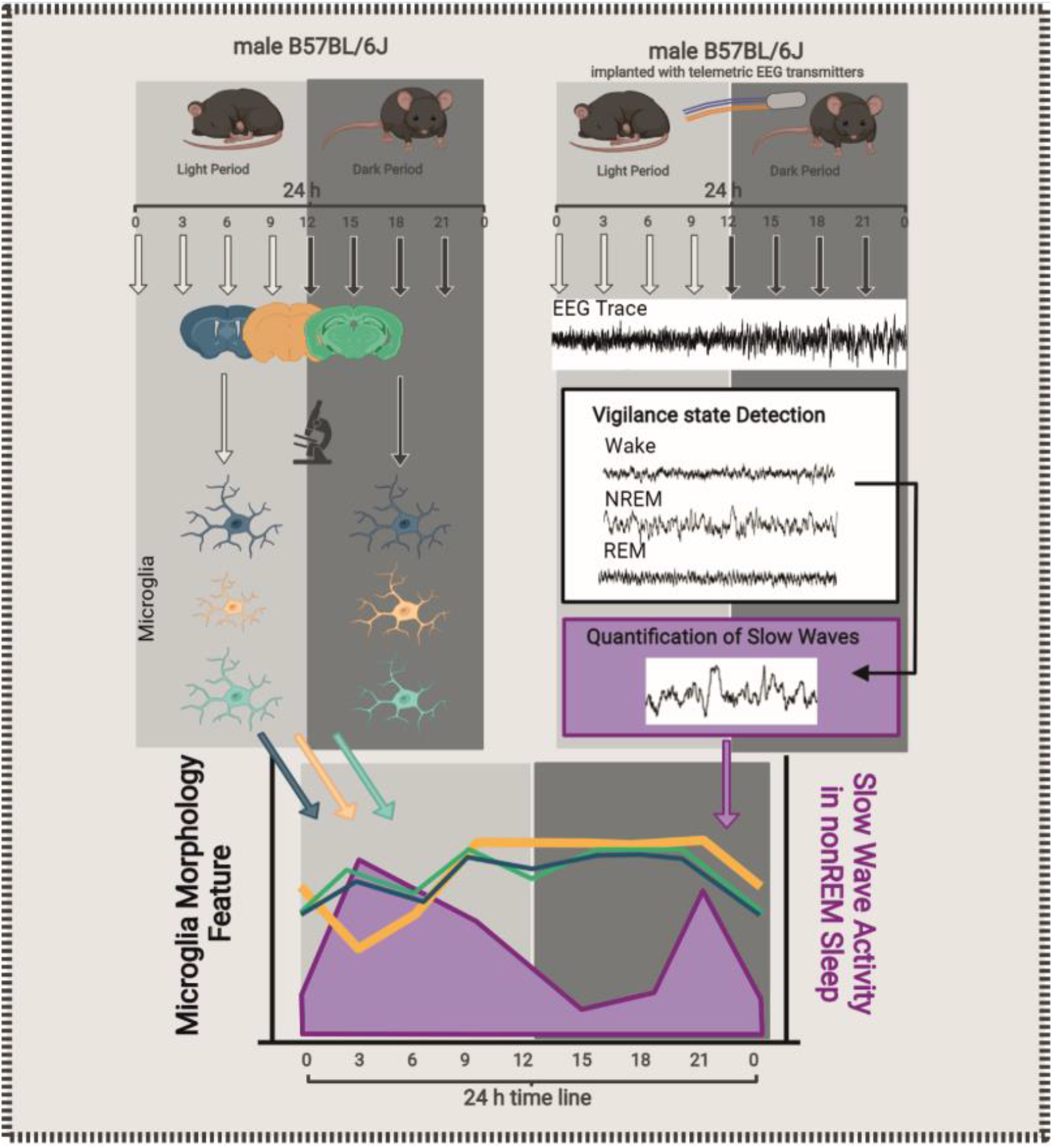

**Main Points:** Microglia show morphological differences between the somatosensory cortex (SC), hippocampus (HC) and basal forebrain (BF) under physiological conditions.

Cortical-, but not HC- or BF microglial cell volume negatively correlates with non-REM sleep slow wave activity.

## Introduction

Sleep deprivation (SD) promotes inflammation in both the periphery and the brain (Mullington et al., 2010). Previous studies reported that long-term total SD induces de-ramification, expression of cytokines and phagocytosis in cortical- and hippocampal microglia (Bellesi et al., 2017; Hsu et al., 2003; Nakanishi et al., 2021; Tuan & Lee, 2019; Wadhwa et al., 2017, 2018; Wisor et al., 2011), the brain’s resident immune cells. These changes closely resemble the microglial response to the gram-negative bacterial cell wall component lipopolysaccharide (LPS), a widely used experimental tool to induce inflammation (Buttini et al., 1996). In contrast, chronic sleep restriction studies have yielded more variable results depending on the species, exposure times, microglial features quantified, and brain areas investigated (Hall et al., 2020; Xie et al., 2020). Recent studies (Corsi et al., 2021; H. Liu et al., 2021) found evidence that microglia also modify vigilance-states, but only few studies (Griffin et al., 2019; Hayashi et al., 2013; Nakanishi et al., 2021) have investigated microglial biology throughout the diurnal inactivity/activity pattern or the natural sleep-wake cycle, both of which are under circadian control (Kuhlman et al., 2018). Moreover, while it is now well known that microglia demonstrate regional heterogeneity (Ayata et al., 2018; Grabert et al., 2016; Stratoulias et al., 2019; Tan et al., 2020), and that they closely monitor the local neuronal activity (Badimon et al., 2020; Cserép et al., 2020; Eyo & Wu, 2013; Li et al., 2012; Ronzano et al., 2021; Tremblay et al., 2010; Umpierre & Wu, 2020; Wake et al., 2009), studies on microglial rhythmicity in sub-cortical areas with diverse neuronal activity patterns, are sparse.

As nocturnal animals, mice are mainly awake during the dark period and sleep mostly during the light period (Franken et al., 1998). Sleep, however, is not uniform throughout the light period. The first hours are dominated by slow (0.5-4 Hz) wave activity. Slow wave activity (SWA) is an electroencephalic (EEG) marker of non-rapid-eye-movement sleep (NREMS) intensity, which reaches its maximum at the beginning of the light period, followed by a well-documented decay (Borbely, 1982; Borbély et al., 2016). During the rest of the light period REM sleep (REMS), lighter NREMS, as well as an increasing number of consolidated waking bouts, prevail.

In the present study, we carefully studied murine microglial morphology as a readout of their function (Beynon & Walker, 2012; Davis et al., 1994; Stence et al., 2001) in physiological conditions throughout the 24h light-dark cycle in three anatomically separate but functionally interconnected brain areas: the somatosensory cortex (SC), the dorsal hippocampus (HC) and the basal forebrain (BF). We investigated whether we could find the previously reported diurnal changes in cortical microglia ramification (Hayashi et al., 2013; Nakanishi et al., 2021) also in the HC or the BF. We then further characterized the physiological alterations in microglial morphology by comparing them to the reactive microglia experimentally induced by the LPS.

Furthermore, to temporally link microglial morphology to vigilance states we performed rodent polysomnography with epidural cortical EEG recordings. The EEG signal arises from the synchronized neuronal population activity (Buzsáki et al., 2012), and the vigilance-state specific neuronal activity patterns are thoroughly mapped for each of the selected brain areas (reviewed in (Adamantidis et al., 2019; Buzsáki, 2015; Holst & Landolt, 2018; Jones, 2020). We were particularly interested in the BF, because it is a key sub-cortical structure regulating both the vigilance states (Xu et al., 2015) and the cortical oscillations (Anaclet et al., 2015).

## Methods

### Animals

All experiments were approved by the Regional State Administration Agency for Southern Finland and conducted in accordance with the Finnish Act on the Protection of Animals Used for Science or Educational Purposes (497/2013). We used 86 (of which 47 went successfully through the entire protocol) male C57BL/6JRccHsd (B6) mice (Envigo, Netherlands) of the age of 10±2 weeks and 26±3 g body weight group-housed in controlled temperature (22 ± 2 °C) with a 12 h light-dark cycle (lights on at 8:00 am, intensity 100 lx) and *ad libitum* access to food and water.

In the lipopolysaccharide (LPS) group (8 animals), the mice were injected intraperitoneally with 5 mg/kg as in Verdonk 2016 (Verdonk et al., 2016) of LPS from *Escherichia coli* (Sigma-Aldrich) dissolved in 0.9 % saline and the control group (8 animals) with 0.9 % saline at *zeitgeber* time point (ZT)6 and perfused 24h later.

### Tissue preparation and immunohistochemistry

To capture the diurnal rhythmicity tissue samples were collected every 3 h in a total time frame of 24 h (64 animals in total; see Fig. 2A).

**Figure 1:**
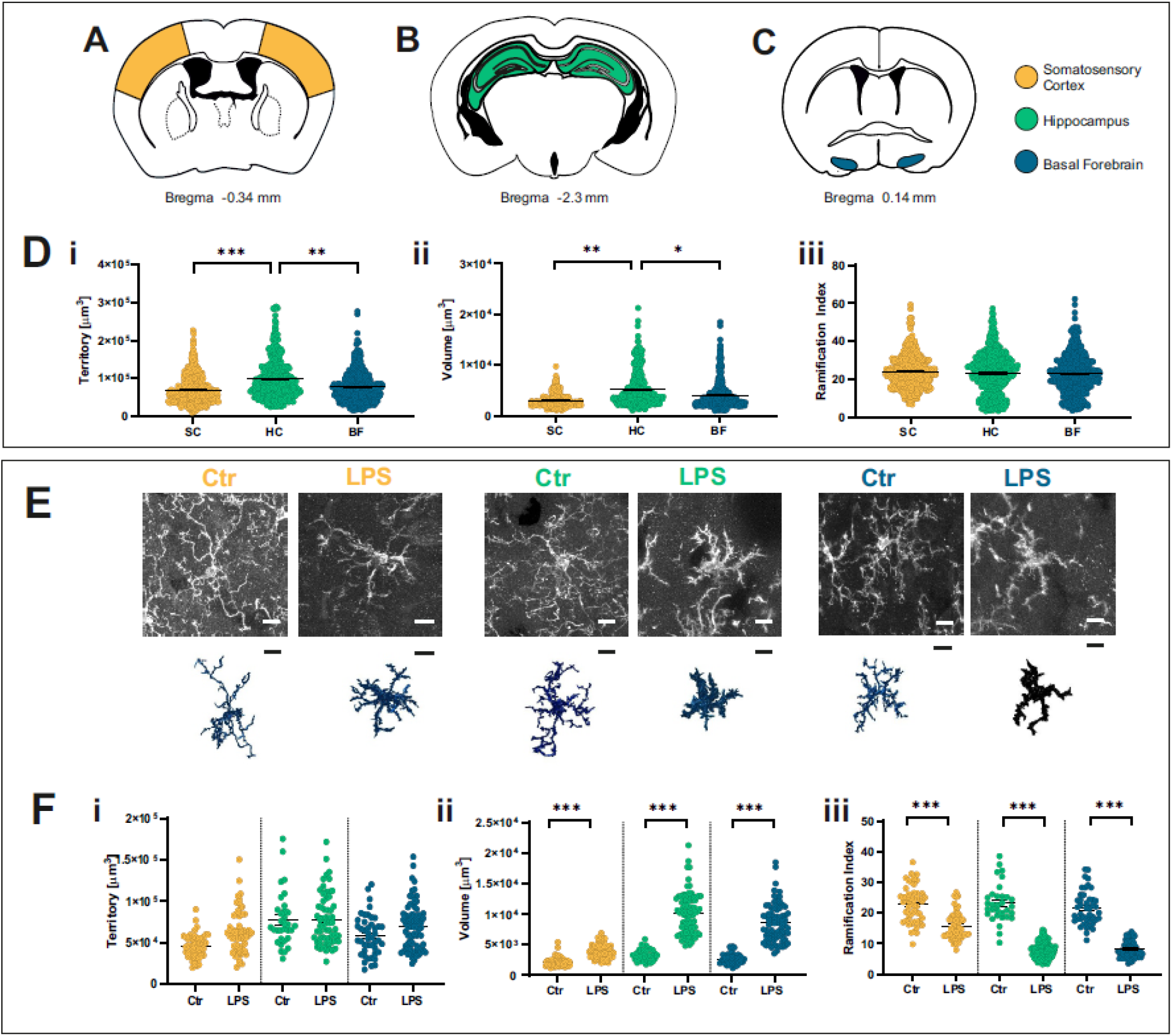
Differences in microglia morphology between brain areas. Microglia morphological differences in somatosensory cortex (SC, yellow) (**A**), hippocampus (HC, green) (**B**) and the basal forebrain (BF, blue) (**C**) under physiological conditions, measured as cellular territory (**Di**) and volume (**Dii**) and ramification index (**Diii**). SC: animals=38; cells=465 HC: animals=29; cells=350 BF: animals=43; cells=386 **E**: Representative confocal z-stack maximum projections and the 3D-reconstructions of microglia in the SC, HC and BF 24 h after a saline (Ctr) injection or a lipopolysaccharide (LPS) injection (5mg/kg, i.p). Tissues collected at ZT *(zeitgeber* time point)6. Scale bars = 10 μm. **F**: LPS injection increases microglia volume and decreases ramification compared to Ctr SC: LPS animals=6 and cells=60; Ctr animals=3 and cells=50 HC: LPS animals=4 and cells=91; Ctr animals=4 and cells=32 BF: LPS animals=4 and cells=74; Ctr animals=5 and cells=44 Statistical significance was assessed with General Estimating Equation (GEE); post hoc Bonferroni. Shown are the means ± SEM; *, *p*<0.05; **, *p*<0.01; ***, *p*<0.001

**Figure 2:**
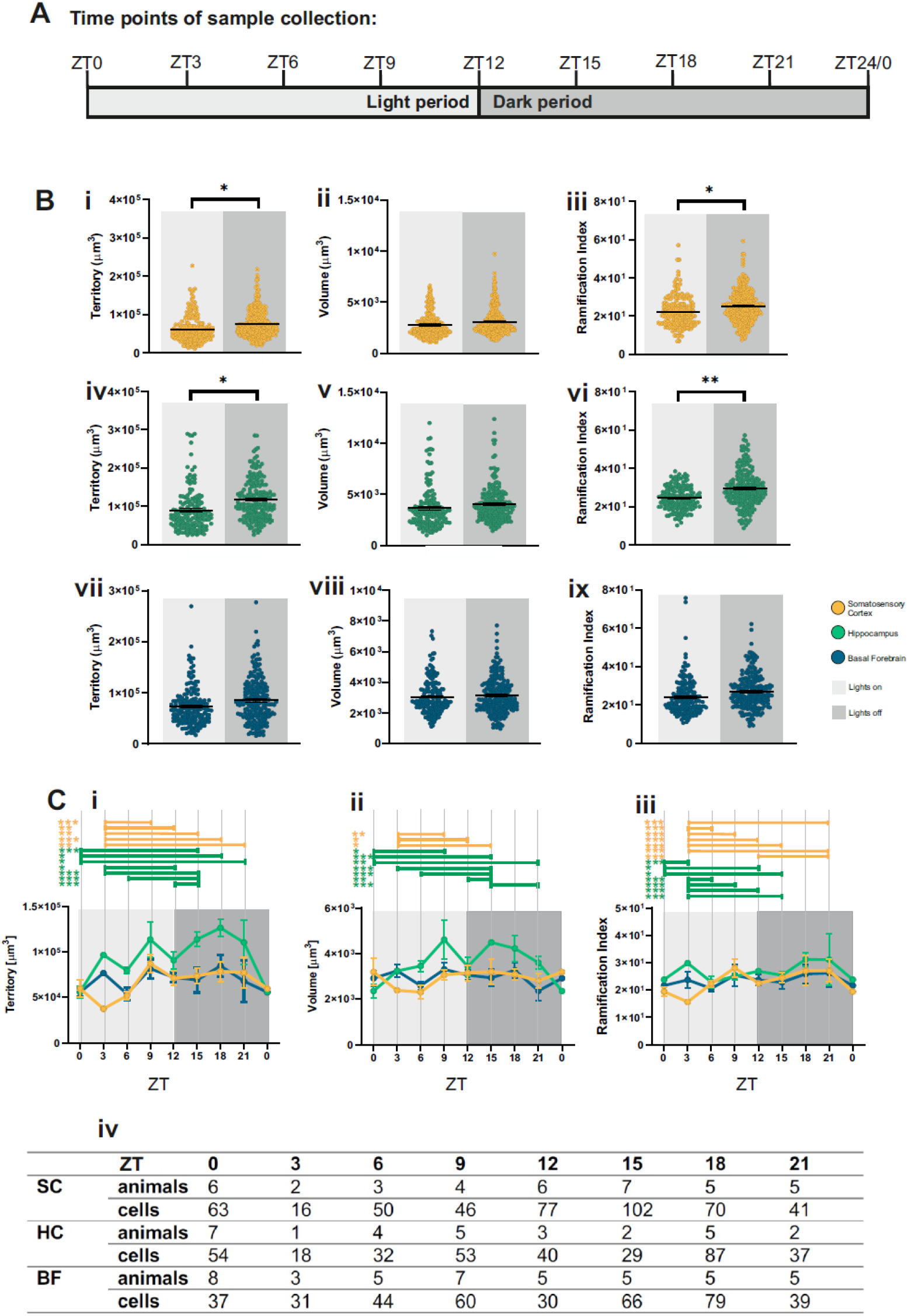
Alterations in microglia morphology between the light and dark period. **A:** Timeline of sample collection during light (12 h; light grey) and dark (12 h; dark grey) period. **B:** Differences in microglia size and ramification complexity between light (ZT0 to ZT9) and dark (ZT12 to ZT21) period can be detected in the SC and HC, but not in the BF. SC: light animals=15 and cells=175; dark animals=22 and cells=290 HC: light animals=17 and cells=157; dark animals=12 and cells=193 BF: light animals=23 and cells=172; dark animals=30 and cells=214 **C:** Alterations in morphological features in 3 h intervals over 24 h, bars indicating statistically significant differences between time points within one brain area. Number of animals/cells in (**C iv**). Displayed are means ± SEM; *, *p*<0.05; **, *p*<0.01; ***, *p*<0.001; Statistical significance was assessed with General Estimating Equation (GEE); post hoc Bonferroni.

The mice were injected with a lethal dose of pentobarbital (120 mg / kg body weight, Mebunat Vet, Orion Pharma, Finland), transcardially perfused with 4 % paraformaldehyde (PFA, 37 % formaldehyde diluted in phosphate buffered saline (PBS, 0.1 M, pH 7.4)) and the brains were post-fixed in 4% PFA for 24h, followed by 48 h cryoprotection by dehydration in 30% sucrose and stored at −80°C.

Coronal sections (35 μm; minimizing compromise of cellular structures while still allowing for complete antibody penetration) of the somatosensory cortex (SC), the dorsal hippocampus (HC) and the basal forebrain (BF) (approximate distance from Bregma, −0.34 mm; −2.30 mm and 0.14 mm, respectively, see Fig. 1 A) were sliced continuously using a Leica CM3050 S cryostat and collected into tris-buffered saline with Tween (TBST; pH 7.2; 0.01 M with 0.05% Triton X).

Free-floating sections were blocked against non-specific binding with 10 % goat serum (NGS, Jackson ImmunoResearch, USA) in TBS-T for 90 min, incubated first with the primary antibody (microglia/macrophage specific polyclonal rabbit anti-IBA1 antibody 1:2000, 234 003, Synaptic Systems, Göttingen, Germany) at 4 °C for 18 h and subsequently with the secondary antibody (AlexaFluor 568 goat-anti-rabbit 1:500; SySy, Germany) for 2 h in room temperature. The bain slices were then mounted on microscope slides (Menzel, 76 x 26 mm) and coverslipped with ImmuMount (both Thermo Fisher Scientific, Finland) and air-dried.

### Image Acquisition

Images (n=5 per brain area per mouse) were acquired using a SP8 confocal microscope (LEICA, Germany) with a 63x oil immersion objective (PL APO CS2) using the z-stack function. The distance between the images within a stack (0.2 μm) was established based on the size of microglial cells and prior trials and aimed to acquire cellular profiles of sufficient resolution for the subsequent morphological analysis. For each microscopic field selected at random as long as at least one microglial cell was present, a z-stack was obtained from 55 to 80 images taken along the z-axis with a scan speed of 400, laser intensity of 4-8 % and a line average of 3.

### Polysomnography with telemetric EEG/EMG recording

A second group of male B6 mice (6 animals) received Carprofen (5 mg/kg; i.p.; FaunaPharma, Finland) as a preoperative analgesic, and a local treatment at the incision site of Lidocaine (Orion, Finland). Under general anesthesia (isoflurane (Attane Vet, Netherlands): 4 % for induction, 1.5-2 % for maintenance) the mice were placed in a stereotaxic device (Kopf Instruments) and implanted with telemetric transmitters (HDX02, Data Sciences International, St. Paul, MN). The operation was adapted from (Papazoglou et al., 2016). For electroencephalography (EEG) recordings 2 biopotential leads were bilaterally placed in the epidural space of the frontal (0.2 mm lateral to midline; 0.8 mm anterior from Bregma) and the parietal area (1.5 mm lateral to midline; 3.4 mm posterior from Bregma) and fixed to the skull with resin-reinforced glass ionomer dental cement (GC FujiCEM 2, Plandent, Finland). The two electromyography (EMG) leads were placed in the cervical trapezius muscle, approximately 1-2 mm apart along the same bundle of muscle fibers with the help of a 20G syringe tip (BD Microlance, Thermo Fisher Scientific). After the dental cement hardened, the skin incision was closed using silk sutures (3-0 coated Vicryl Plus; Ethicon; USA). Subsequently, the mice were carefully monitored for post-operative recovery. Ten days after the surgery the recording (sampling rate: 500 Hz; bandwidth: 0.5-80 Hz) was started. The recording system (all Data Sciences International) consisted of receiver plates (PhysioTel Receiver, model RPC-1) placed beneath the cages to receive the telemetrically transmitted data from the implants, and a DSI Matrix 2.0 (MX2) managing the communication between the implants, the receiver plates and the acquisition software. The data was recorded with the DSI Talker interface and Spike2 (Version 8.17; Cambridge Electronic Design (CED), England). To avoid a microglial reaction to the EEG electrode implantation, we measured the vigilance states in separate animals.

### Data Analysis

#### Image Analysis

Microglial cells were traced and analyzed with the 3Dmorph (York et al., 2018), a MATLAB-based script that quantifies microglial morphology from 3D data in a semi-automatic and unbiased way.

First, the images were pre-processes by despeckling and enhancing the contrast (0.3-3 %) with Fiji (ImageJ, NIH, USA). Subsequently, the image background noise was reduced, merged cells were separated and the smallest (debris or fragmented branches) and biggest (merged cells that could not be separated due to overlapping branches) objects were manually defined and excluded with the help of 3DMorph Matlab script interfaces. The 3DMorph Matlab script then skeletonized the cells, and traced and identified their branch- and endpoints automatically.

We measured the following features: cell territory (volume of a 3D polygon around the cell’s external points), cell volume (determined by the number of voxels multiplied by the scale to convert into real world units), and ramification index (ratio of the cell’s territory to its volume).

Chosen for its outstanding non-biased automatic and very detailed quantification of microglia features, the 3DMorph method proved as sensitive to minor quality differences in images, leading to the exclusion of images/animals: LPS + saline original 8+8 reduced to 6+5; diurnal rhythmicity samples original 8×8=64 reduced to 47 animals (for more details see Figure 2 C iv).

Image processing and cell tracing were performed by two investigators blind to the experimental condition.

#### Sleep scoring and calculation of SWA

All EEG data was divided into 4 s epochs with a Spike2 manual sleep scoring script (sleepscore_v1.01; CED) and defined as wake, NREMS, REMS or noise based on the standard criteria and as described previously (Mäkelä et al., 2010). In short: Wakefulness was identified as low-amplitude high-frequency (>10 Hz) EEG activity combined with high EMG activity; NREMS was scored as high-amplitude EEG delta (0.5–4 Hz) waves in combination with low EMG activity; and REMS was characterized by low-amplitude EEG theta (5–9 Hz) activity with minimal or absent EMG activity. Artifacts were scored as noise and excluded from further analysis.

EEG power spectra were calculated within the 0.5-30 Hz frequency range by fast Fourier transform (FFT = 1024, Hanning window, 0.5 Hz resolution) in 3h-time bins for NREMS in Spike2. Before further analysis, all EEG spectra were normalized to the total power of the recording day for each individual animal to control for the non-specific effects of recording instrumentation. From the power spectrum we focused on the SWA i.e. delta (0.5-4 Hz) power band during NREMS epochs in 3h-bins. We also calculated cumulated delta power as in (Stenberg et al., 2003) by multiplying the delta power with the number of epochs in 3h-bins (see Supplementary Table 1).

### Statistical Analysis

Statistical analyses were performed using Graphpad Prism (v. 8.2.0, SanDiego, USA) and SPSS version 25 (IBM Corp., IL, USA).

#### GEE and correction for multiple testing with Bonferroni

Brain area differences, the effects of the LPS treatment, differences between the light and the dark period and the time point differences were assessed using generalized estimating equations (GEEs). This method controls for partial within-subject dependencies, a crucial feature for our data set since we analyzed microglia cells grouped by sampling time points and brain area, and not grouped by animals. To ensure statistical power despite the varying group sizes, data sets were tested for normality and variance homology.

Multiple testing correction was performed by a planned post hoc Bonferroni method. Results were expressed as mean ± SEM, and p values < 0.05 were considered statistically significant.

#### Data randomization for Spearman correlation

For the Spearman correlation analysis we needed to obtain an equal number of measures for cumulated delta power (4 light period time-points derived from 6 animals) and for morphological features (4 light period time points derived from differing number of animals and cells, see Figure 2 C iv). We accomplished this by pooling morphological feature measurements by assigning each microglial cell a random number and established six groups by following the numbers in ascending order within the same ZT time point. The results were tested against a Bonferroni-adjusted alpha level of 0.016 (0.05/3).

## Results

We investigated murine microglial morphology throughout the 24 h light-dark cycle in the somatosensory cortex (SC), dorsal hippocampus (HC) and the basal forebrain (BF). Microglia morphology was quantified using 3D-cell-reconstructions from z-stack confocal images of IBA1-immunostained microglia in SC, HC and BF of male B6 mice (Fig. 1A-C and 1E). Microglia territory and volume were used to define the cell size, and the ramification index as measure for ramification complexity. We found differences in microglial morphology between: 1) the brain areas, 2) the dark-and light periods, 3) the vigilance states 4) and specific sleep features (SWA).

### Differences in microglia morphology between brain areas

To analyze region dependent differences, we compared the samples collected from SC, HC and BF. Microglia size in the HC differed significantly from both the SC and BF. More precisely, cell territory (GEE with post hoc Bonferroni; SC vs. HC: *p*=2.3xE^-6; HC vs. BF: *p*=2xE-3) and cell volume (GEE; post hoc Bonferroni; SC vs. HC: *p*=7xE-3; HC vs. BF: *p*=0.032) were smaller in the SC and BF as compared to the HC (Fig. 1 A-D).

In response to LPS injection, microglia in all investigated brain areas were similarly affected. In accordance with previous results, we observed that 24 h after the LPS injection, the cell volume in all brain areas increased (GEE; post hoc Bonferroni; SC: *p*=7.64xE-6; HC: *p*=3.0xE-15;BF: *p*=1.14xE-7), whereas the ramification index decreased (GEE; post hoc Bonferroni; SC: *p*=1.44xE-11; HC: *p*<1xE-15;BF: *p*=2.01xE-5) (Fig. 1 E-F).

### Differences in microglia morphology between the light and dark period

To investigate the differences between light and dark period, we divided the samples into two groups: The light period consisted of samples taken at ZT0, ZT3, ZT6 and ZT9, and the dark period of ZT12, ZT15, ZT18 and ZT21 (Fig. 2 A).

We found differences in SC and HC, but not in BF (Fig. 2 B).

The cell territories in SC and HC (GEE; post hoc Bonferroni; SC: *p*=3.6xE-3; HC: *p*=3.3xE-3) and the ramification index in the HC (GEE; post hoc Bonferroni; *p*=1.4xE-3) were larger during the dark period as compared to the light period.

### Alterations in microglia morphology within the light and dark periods

To address the microglial morphology dynamics within the light and dark period more closely, we compared the sampled time points (3 h intervals) separately (Fig. 2 C). Within the dark period, no clear pattern emerged, whereas within the light period we found statistically significant changes in SC and HC, but not in the BF (GEE; post hoc Bonferroni; Fig. 2 Ci-ii).

More precisely, during the light period, microglia territory and ramification index in the SC decreased until reaching a minimum at ZT0/ZT3 before peaking at ZT9 (Fig. 2 Ci). Microglia in the HC exhibited seemingly more random fluctuations within the light period, which were for territory and ramification index overall lower than the average of the dark period (Fig. 2 Ci-ii).

### Alterations of microglia morphology in SC coincide with NREMS SWA

Most of the statistically significant differences in microglial morphology occurred during the light period as compared to the dark (Fig. 2 C and Fig. 3 B). We therefore investigated the light period time points by comparing them to the microglia feature- and brain area-specific average of the dark period as a reference (Fig. 3 B; dashed line). Since the light period is the habitual sleep period for mice, we hypothesized that the sleep-wake cycle within the light period is temporally associated with microglial morphology. We therefore performed polysomnography in a separate group of animals (Supplementary Table 1.).

**Figure 3:**
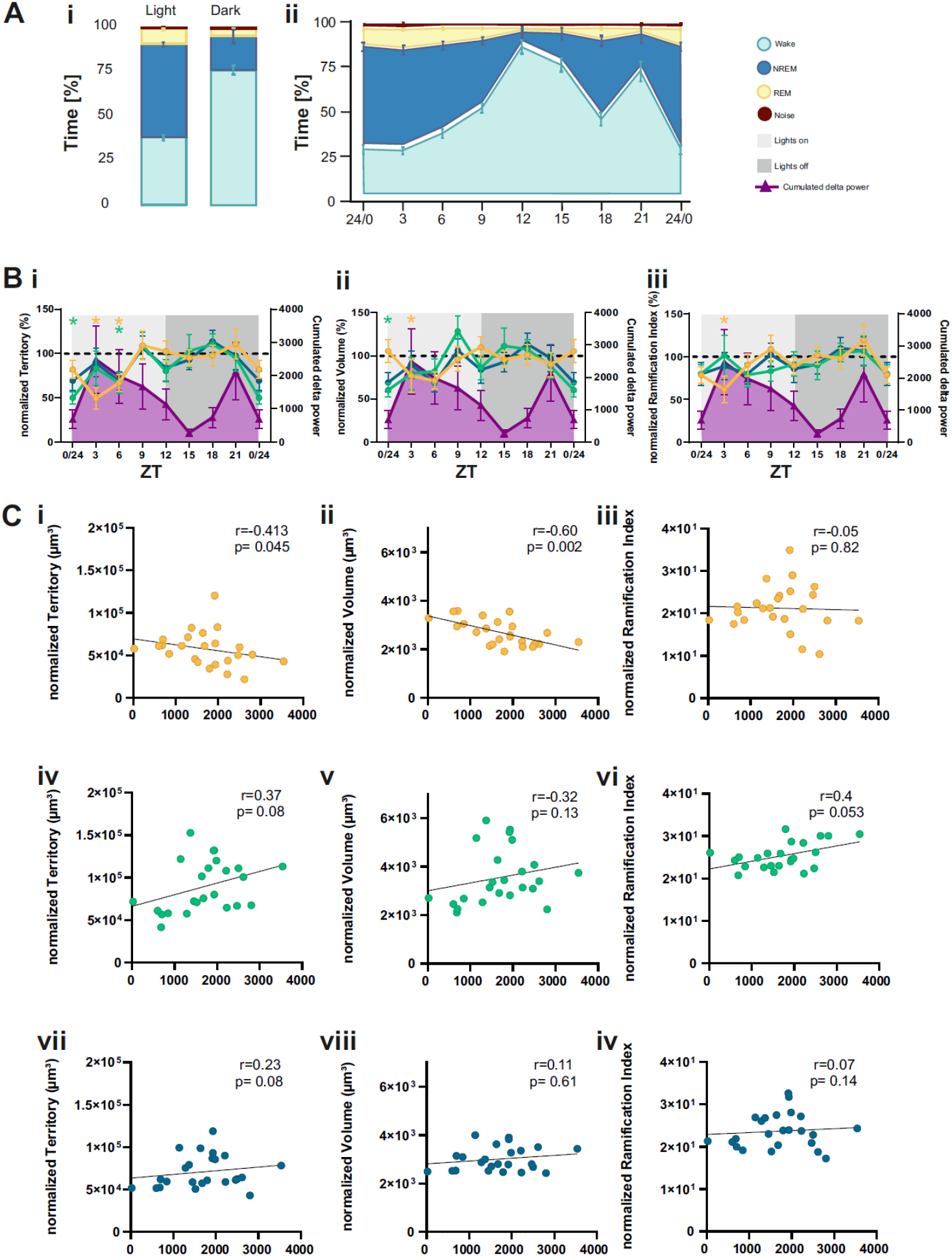
Alterations in microglia morphology between vigilance states coincide with NREMS SWA. **A:** Vigilance state distribution during the 12h light (left) and the 12h dark (right) period (**Ai**) and throughout 24h in 3h-resolution (**Aii**). Vigilance states were scored as wake (light green), NREMS (blue) and REMS (yellow), and periods of noise. Animals=6 **B:** Normalized microglia features (territory (**i**); volume (**ii**); ramification index (**iii**)) and cumulated delta power in NREMS (violet) in 3h time bins over 24h. Significant differences from the baseline (mean of all dark period measurements, dashed black line) are marked with asterisks, color-coded for the respective brain area. Cumulated delta animals=6; for n animals/cells of morphology features, see Fig. 2 C **C:** Spearman correlation between cumulated delta power during NREMS (4 light period time-points derived from 6 animals) and microglia morphology feature measurements (4 light period time points derived from differing number of animals and cells, see Figure 2 C iv, pooled into groups within equal ZTs) in 3h-time bins during the light period split by brain area. The results were tested against a Bonferroni-adjusted alpha level of 0.016.

During the first six hours of the light period cell territory (GEE; post hoc Bonferroni; ZT0: HC *p*=5.36xE-12; ZT3: SC: *p*<1xE-15; ZT6: SC *p*=2.27xE-3; HC *p*=2.13xE-6) and volume (GEE; post hoc Bonferroni; ZT0: HC *p*=1.6xE-3; ZT3: SC: 8.7xE-3) in SC and HC, and ramification index in the SC (GEE; post hoc Bonferroni; ZT3 SC: *p*=2.16xE-11) and were significantly lower compared to the dark period reference. This occurred in parallel with intense NREMS with maximal SWA, as quantified by multiplying the number of NREMS epochs with delta power (cumulated delta power) (Fig. 3 B).

To investigate this relationship between microglia morphology and SWA further, we performed a correlation analysis which showed that microglial cell volume in the SC negatively correlated with cumulated delta power (Spearman correlation with Bonferroni-adjusted alpha level of 0.016; SC volume: r=-0.6, *p*=2.0xE-3; Fig. 3 Ci-iii). HC and BF did not show a statistically significant correlation (Fig. 3Civ-ix).

## Discussion

### Diurnal alterations in microglial morphology were brain region specific

We quantified microglial morphology by utilizing a 3D reconstruction method (York et al., 2018), in three functionally different brain areas of mice throughout the 24 h light-dark cycle.

As reported before (Grabert et al., 2016; Stratoulias et al., 2019) microglia demonstrated regional heterogeneity in their features (Fig. 1 D), however the stimulation with LPS evoked a comparable response in all brain regions indicated by volume increase and ramification complexity decrease (Fig. 1 E and 1 F). The combination of morphological changes and the magnitude in which they change under the influence of LPS differed from the physiological changes we observed over a 24h period (compare Figure 1 E-F with Figure 2 B).

Brain area heterogeneity was evident between 12h light and 12h dark period (Fig. 2B). Microglial ramification in the SC and HC was higher in the 12 h dark period as compared to the 12 h light period; a finding in line with previous studies reporting cortical microglial hyper-ramification during the dark period and de-ramification during the light period (Hayashi et al., 2013; Nakanishi et al., 2021). Notably, microglia in the BF did not demonstrate morphological differences between the light- and the dark period (Fig. 2B vii-ix). As circadian regulation is both cell-autonomous and centrally orchestrated (reviewed in (Mohawk et al., 2012)) it should be evident in all brain areas, including in the BF (Nikonova et al., 2017). Our findings argue against the hypothesis that circadian regulation either directly via local expression of circadian genes and/or in combination with the superchiasmatic nucleus (SCN) mediated orchestration, would exclusively be responsible for the reported changes in microglial morphology. Furthermore, the time course of morphological alterations in the cortex did not follow a typical 12 h light to 12 h dark diurnal pattern with opposing characteristics in the dark-as compared to light period. Instead, the majority of the statistically significant alterations in microglial morphology (Fig. 2C, upper panel) took place within the light period (the habitual sleeping period of the mice). In contrast, during the dark period (the murine active period) microglial morphology remained relatively stable in all brain areas.

As microglia closely monitor neuronal activity (Badimon et al., 2020; Cserép et al., 2020; Eyo & Wu, 2013; Li et al., 2012; Ronzano et al., 2021; Tremblay et al., 2010; Umpierre & Wu, 2020; Wake et al., 2009), and as the vigilance-state specific regional neuronal activity patterns are well characterized (reviewed in (Adamantidis et al., 2019; Holst & Landolt, 2018) for all the selected brain regions (Jones, 2017), we propose that these regional differences in neuronal activity could explain our findings. Under physiological conditions, microglial morphology is mainly regulated by the local neuronal activity associated changes in extracellular ionic--and ATP concentrations (Reviewed in (Izquierdo et al., 2019)).

### Microglial morphology during the second half of the light period and throughout the dark period did not differ between the brain areas

During the second half of the light period REMS, light NREMS (as quantified by the decreased SWA), and an increasing number of wake bouts prevailed whereas during the entire dark period the mice were mostly awake (Fig 3 A, Supplementary Table 1)(Franken et al., 1998). Throughout this time, microglial morphology did not show major alterations nor differ between the brain areas (Fig. 3 B). We propose that this can be explained by the vigilance-state specific neuronal activity patterns in the investigated brain areas.

In waking and in REMS when the EEG displays continuous high frequency activity, both the cortical- and hippocampal networks switch into a continuous fast firing mode (Adamantidis et al., 2019; Holst & Landolt, 2018). The rodent hippocampal neurons also enter into a prominent theta (5-9 Hz) oscillation with nested high frequency gamma (30-70 Hz) oscillations (Buzsáki, 2015; Vanderwolf, 1969). In the sub-cortical BF, which is composed of functionally diverse groups of neurons, the most numerous groups are the wake- and REMS active neurons discharging in association with fast cortical activity (Jones, 2017). Taken together, in all investigated brain areas, neuronal activity in waking and REMS resemble each other (i.e. continuous high frequency firing activity in association with fast EEG oscillations), and could therefore explain why microglia morphology did not differ between the brain areas during the second half of the light period and throughout the entire dark period. Nonetheless, it needs more studies to investigate the mechanisms behind this connection.

### Vigilance state-specific neuronal activity may influence microglial morphology in a brain-site specific manner

During the first 3 hours of the light period, when intense NREMS with maximal SWA dominate (Fig 2 A+B, Supplementary Table 1) the cortical microglia displayed minimum ramification complexity, reversing back to the dark period level by the end of the light period (Fig. 2 C). This pattern coincides with maximal NREMS SWA and its decay (Fig 3 B). We found that cortical microglial territory and volume negatively correlated with NREMS SWA, while the microglial features in the HC or the BF did not (3C). Even though further studies are needed to find the mechanisms behind the correlation, we propose that vigilance-state specific neuronal activity could explain this connection.

In the cortex, the neuronal correlates of cortical biphasic SWA during NREMS are well characterized: The bursting activity (UP-STATE) of the depolarized cortical neurons is periodically (0.5-4.0 s) intersected with hyperpolarized silent phases (DOWN-STATES) (Steriade et al., 1993; Steriade & Timofeev, 2003). Simultaneously, the hippocampal neuronal ensembles engage into re-occurring sharp-wave-ripple (110-200 Hz) oscillations, which are cross-frequency coupled to cortical SWA. According to several reports (Badimon et al., 2020; Cserép et al., 2020; Eyo & Wu, 2013; Li et al., 2012; Ronzano et al., 2021; Tremblay et al., 2010; Umpierre & Wu, 2020; Wake et al., 2009), microglia are sensitive to the fluctuations in overall neuronal activity rather than the type (i.e. depolarization/hyperpolarization) of activity causing it. In other words, if the neuronal signal is predictable, it is less likely to affect microglia. Interestingly, a recent human study demonstrated that the complexity of neuronal activity during sleep SWA was significantly lower as compared to both wakefulness and REMS (Stevner et al., 2019). We propose that neuronal activity during the biphasic cortical SWA with its predictable and recurring activity pattern, would not stimulate microglial ramification. In contrast, in the HC, where the neurons engage into more irregular, high frequency ripple oscillations during NREMS, microglial ramification complexity would remain as stimulated as in waking. Similarly, in the BF, when the wake- and REMS active cells decrease their firing rate during NREMS SWA, another group of cells, coupled to cortical SWA, increases it (Jones, 2017). Hence, a mere redistribution of neuronal firing would continue to stimulate BF microglial ramification throughout the light period.

### The neuromodulatory milieu may influence microglial morphology in a brain-site specific manner

In association with neuronal activity, the neuromodulatory milieu varies across vigilance states and influences microglial morphology (reviewed in (Izquierdo et al., 2019)). Recently, two *in vivo*-imaging studies (Y. U. Liu et al., 2019; Stowell et al., 2019) reported that norepinephrine (NE) inhibits cortical microglial dynamic contacts to synapses in waking, when NE levels are high, and demonstrated a release of inhibition under anesthesia or NE agonist treatment. These studies, however, did not directly study NREMS, when monoamine release naturally decreases or REMS when monoamine release is minimal. While anesthesia can be a useful tool to investigate restricted aspects of sleep, the neuronal activity patterns and particularly the temporal dynamics of sleep dependent neuronal activity are lost (for review see for example (Franks & Wisden, 2021)).

In addition to neurotransmitters, the cortical extracellular ionic composition fluctuates in association with SWA (Ding et al., 2016). In transition from waking to NREM SWA, both [Ca^2+^]_ext_ and [Mg^2+^]_ext_ decrease while [K^+^]_ext_ levels decrease. Microglial ramification (and their surveillance) in physiological conditions critically depends on resting membrane potential, and the tonic activity of THIK-1 K+ channels (TWIK-related Halothane-Inhibited K+ channel) (Madry, Kyrargyri, et al., 2018). It is possible that the vigilance state–specific cortical changes in the extracellular ionic composition contribute to the negative correlation with microglial ramification complexity and SWA. The most widely studied regulator of microglial morphology is P2Y12 receptor mediated ATP chemotaxis (Ayata et al., 2018; Davalos et al., 2005; Haynes et al., 2006; Nimmerjahn et al., 2005). Neuronal activation locally releases ATP from both astrocytes (Pascual et al., 2005) and neurons (Pankratov et al., 2006), contributing to ramification, and chemotaxis of microglial processes towards ATP (Honda et al., 2001). Simultaneously, ATP/ADP is converted into AMP via the ectoenzyme CD39 (NTPDase1) and then further to adenosine by CD73 (5′-nucleotidase), which together with the equilibrate nucleotide transmitter (ENT1) contribute to elevate the local extracellular adenosine concentration. Adenosine in turn potentiates both spontaneous- and ATP-dependent microglial ramification (Madry, Arancibia-Cárcamo, et al., 2018; Matyash et al., 2017; Wollmer et al., 2001). In general, the extracellular levels of adenosine, and ATP, closely correlate with neuronal activity (Dunwiddie & Masino, 2001) and are higher in waking than in sleep, but with distinct regional differences (Dworak et al., 2010; T. Porkka-Heiskanen et al., 2000; Tarja Porkka-Heiskanen et al., 1997). We propose that together with the aforementioned factors, the local differences in the extracellular ATP/ADE could explain why the BF microglia remain stably ramified throughout the 24-h period. Dvorak et al., 2010 demonstrated that the extracellular levels of ATP in mice are much higher in the BF and frontal CX, as compared to cingulate cortex or hippocampus during the first 3h of the light period. It also took longer for the BF ATP levels as compared to other brain areas to return to the dark period levels, thus providing a possible explanation for the stability of the ramified microglial phenotype in the BF.

In conclusion, in our study we show that under physiological conditions BF microglia differ from cortical- and hippocampal microglia, while after immunostimulation, the responses are similar in all brain areas. In addition, the cortical microglial morphology negatively correlates with sleep SWA, suggesting that microglia are sensitive to local differences in vigilance-state specific neuronal activity.

## Acknowledgements

The authors would like to thank B.Sc. Cara Gebauer for her excellent technical assistance, Dr. Mikaela Laine for strong support with the statistical data analysis and for giving feedback on the manuscript, Prof. Iiris Hovatta for general supervision. Special thanks to Liljeström and the Biomedicum Imaging Unit (BIU), and Marko Crivaro and the Light Microscopy Unit, Helsinki, for kind help and technical assistance with the imaging work. Financial support was provided by the Finish Medical Association Läkaresellskapet, the Signe and Ane Gyllenberg Foundation and the Brain & Mind Doctoral Programme of the University of Helsinki.

The graphical abstract was constructed using BioRender (https://biorender.com/) and the figures with CorelDraw version 2021 (www.coreldraw.com).

