## Supplementary Table for "Physiological alterations in microglial morphology associate with the sleep-wake cycle in a brain region-specific manner"

| ZT / State | | Wake | SEM | SWS | SEM | Norm. delta power | SEM | REM | SEM | Noise | SEM |
|---|---|---|---|---|---|---|---|---|---|---|---|
| 0 / 24 | Epochs | 758.83 | ± 80.25 | 1662.50 | ± 72.91 | 7.13E-08 | ± 1.37E-08 | 256.50 | ± 22.48 | 19.17 | ± 5.46 |
| | % | 28.14 | ± 2.98 | 61.64 | ± 2.70 | | | 9.51 | ± 0.83 | 0.71 | ± 0.20 |
| 3 | Epochs | 738.50 | ± 62.71 | 1623.33 | ± 79.14 | 8.23E-08 | ± 1.84E-08 | 301.33 | ± 21.66 | 33.83 | ± 31.05 |
| | % | 27.38 | ± 2.33 | 60.19 | ± 2.93 | | | 11.17 | ± 0.80 | 1.25 | ± 1.15 |
| 6 | Epochs | 1037.00 | ± 76.08 | 1405.33 | ± 72.96 | 6.77E-08 | ± 1.5E-08 | 249.33 | ± 23.08 | 5.33 | ± 2.70 |
| | % | 38.45 | ± 2.82 | 52.11 | ± 2.71 | | | 9.24 | ± 0.86 | 0.20 | ± 0.10 |
| 9 | Epochs | 1474.83 | ± 80.45 | 1048.67 | ± 67.07 | 6.61E-08 | ± 1.47E-08 | 168.50 | ± 19.25 | 5.00 | ± 3.26 |
| | % | 54.68 | ± 2.98 | 38.88 | ± 2.49 | | | 6.25 | ± 0.71 | 0.19 | ± 0.12 |
| 12 | Epochs | 2532.33 | ± 118.62 | 135.67 | ± 103.62 | 5.91E-08 | ± 1.57E-08 | 23.00 | ± 16.20 | 6.00 | ± 2.74 |
| | % | 93.89 | ± 4.40 | 5.03 | ± 3.84 | | | 0.85 | v0.60 | 0.22 | ± 0.10 |
| 15 | Epochs | 2221.50 | ± 114.90 | 426.83 | ± 101.21 | 0 | 0 | 35.33 | ± 14.19 | 13.33 | ± 6.47 |
| | % | 82.37 | ± 4.26 | 15.83 | ± 3.75 | | | 1.31 | ± 0.53 | 0.49 | ± 0.24 |
| 18 | Epochs | 1284.83 | ± 111.39 | 1237.33 | ± 89.60 | 1.04E-07 | ± 2.65E-08 | 170.67 | ± 26.26 | 4.17 | ± 2.79 |
| | % | 47.64 | ± 4.13 | 45.88 | ± 3.32 | | | 6.33 | ± 0.97 | 0.15 | ± 0.10 |
| 21 | Epochs | 2109.83 | ± 172.16 | 525.50 | ± 152.54 | 9.29E-08 | ± 2.03E-08 | 49.50 | ± 19.42 | 12.17 | ± 6.48 |
| | % | 78.23 | ± 6.38 | 19.48 | ± 5.66 | | | 1.84 | ± 0.72 | 0.45 | ± 0.24 |

*Table 1 Distribution of vigilance states (Wake; SWS = slow wave sleep; REM) and normalized delta power over 24h; mean +/- SEM., for each zeitgeber time point (ZT) in absolute values (epochs = 4s long time bins) and percentage;* $N_{animals}$*=6*
